# Viromic analysis of wastewater input to a river catchment reveals a diverse assemblage of RNA viruses

**DOI:** 10.1101/248203

**Authors:** Evelien M. Adriaenssens, Kata Farkas, Christian Harrison, David L. Jones, Heather E. Allison, Alan J. McCarthy

## Abstract

Detection of viruses in the environment is heavily dependent on PCR-based approaches that require reference sequences for primer design. While this strategy can accurately detect known viruses, it will not find novel genotypes, nor emerging and invasive viral species. In this study, we investigated the use of viromics, i.e. high-throughput sequencing of the biosphere viral fraction, to detect human/animal pathogenic RNA viruses in the Conwy river catchment area in Wales, UK. Using a combination of filtering and nuclease treatment, we extracted the viral fraction from wastewater, estuarine river water and sediment, followed by RNASeq analysis on the Illumina HiSeq platform for the discovery of RNA virus genomes. We found a higher richness of RNA viruses in wastewater samples than in river water and sediment, and assembled a complete norovirus GI.2 genome from wastewater effluent, which was not contemporaneously detected by conventional qRT-PCR. To our knowledge, this is the first environmentally-derived norovirus genome sequence to be available from a public database. The simultaneous presence of diverse rotavirus signatures in wastewater indicated the potential for zoonotic infections in the area and suggested run-off from pig farms as a possible origin of these viruses. Our results show that viromics can be an important tool in the discovery of pathogenic viruses in the environment and can be used to inform and optimize reference-based detection methods provided appropriate and rigorous controls are included.

**Importance:** Enteric viruses cause gastro-intestinal illness and are commonly transmitted through the faecal-oral route. When wastewater is released into river systems, these viruses can contaminate the environment. Our results show that we can use viromics to find the range of potentially pathogenic viruses that are present in the environment and identify prevalent genotypes. The ultimate goal is to trace the fate of these pathogenic viruses from origin to the point where they are a threat to human health, informing reference-based detection methods and water quality management.

## Introduction

Pathogenic viruses in water sources are likely to originate primarily from contamination with sewage. Classic marker bacteria used for faecal contamination monitoring, such as *Escherichia coli* and *Enterococcus* spp., are not, however, good indicators for the presence of human enteric viruses (1). The virus component is often monitored using qPCR approaches, which can give information on the abundance of specific viruses and their genotype, but only those that are both known and characterised (2). Viruses commonly targeted in sewage contamination assays include noroviruses (3), hepatitis viruses (4), enteroviruses (5), and various adenoviruses (6, 7). Viral monitoring in sewage has previously yielded positive results for norovirus, sapovirus, astrovirus, and adenovirus, indicating that people are shedding viruses that are not necessarily detected in a clinical setting (8). This same study found a spike in norovirus genogroup GII sequence signatures in sewage two to three weeks before the outbreak of associated disease was reported in hospitals and nursing homes. The suggestion, therefore, is that environmental viromics can provide an early warning of disease outbreaks, in addition to the monitoring of virus dissemination in watercourses.

Recent reviews have proposed the use of viral metagenomics or viromic approaches as an alternative method to test for the presence of pathogenic viruses in the environment (2, 9, 10). Provided the entire viral community is sampled and sequenced, novel genotypes or even entirely novel viruses can be detected. Potential new viral markers for faecal contamination have already been revealed, such as pepper mild mottle virus and crAssphage (11, 12), among the huge diversity of human viruses found in sludge samples (13).

In this pilot study, we have used viromics to investigate the presence of human pathogenic RNA viruses in wastewater, estuarine surface water and sediment in a single catchment. The water and sediment samples were collected at, and downstream of, the wastewater treatment plant (Lanrwst, Wales, UK), at the estuary of the river Conwy near a bathing water beach (Morfa, Wales, UK) (Figure 1). To our knowledge, this is the first study to use unamplified environmental viral RNA for sequencing library construction, sequence dataset production and subsequent analysis. Because we used a directional library sequencing protocol on RNA, rather than amplifying to cDNA, we were able to distinguish single-stranded from double-stranded RNA genome fragments.

**Figure 1:**
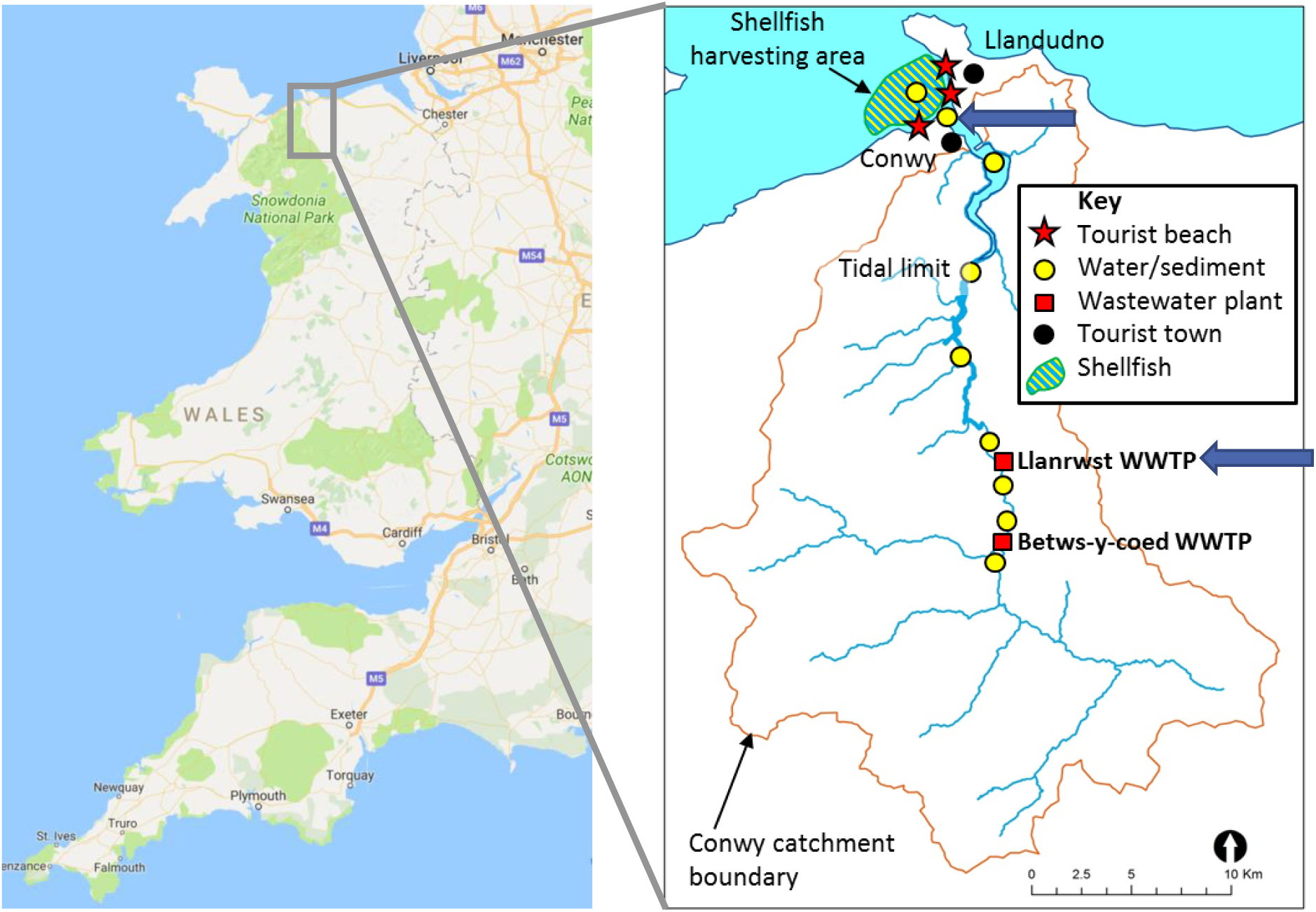
Map of the sampling locations, indicated with blue arrows. Data in the left panel was taken from Google Maps.

## Results

### Sample overview

Wastewater influent and effluent samples were collected from the Llanrwst wastewater treatment plant (53°08’24.4“N 3°48’12.8“W; Figure 1) in September and October 2016, resulting in four different samples, LI_13-9 (Llanrwst influent Sep 2016), LE_13-9 (Llanrwst effluent Sep 2016), LI_11-10 (Llanrwst influent Oct 2016), LE_11-10 (Llanrwst effluent Oct 2016). Estuarine surface water (SW) was collected from Morfa beach (53°17’37.7“N 3°50’22.2“W; Conwy, Wales, Figure 1) in November 2016 and sediment from the same site in October and November 2016 (Sed1, Sed2, respectively).

As an initial assessment, samples were tested for the presence of a subset of locally occurring enteric RNA viruses using qRT-PCR (Table 1). Only norovirus (NoV) genogroup GII signatures were detected in the wastewater samples. In the samples collected in September 2016, 10^3^ genome copies (gc)/l of norovirus GII were observed in both the influent (LI_13-9) and in the effluent (LE_13-9). In the samples collected in October 2016, approx. 10^2^ gc/l (below the limit of quantification which was approx. 200 gc/l) were observed in the influent (LI_11-10) and a considerably higher concentration of 5×10^4^ gc/l was noted in the effluent (LE_11-10). All qRT-PCRs were negative for the presence of sapoviruses (SaV) and hepatitis A/E viruses (HAV/HEV). None of the target enteric viruses were found in the surface water and sediment samples.

**Table 1:** Summary of viromic and qRT-PCR detection of the presence of specific RNA viruses across the samples (sewage, estuarine water and sediment).

| Sample name<sup>a</sup> | Sample volume/mass | Location | # contigs (curated) | Target RNA viruses detected in contigs<sup>b</sup> | qRT-PCR results (gc/l)<sup>c</sup> |
| --- | --- | --- | --- | --- | --- |
| LI_13-9 | 1 l | Llanrwst WWTP | 5721 | RVA, RVC, PBV, SaV | NoVGII (1,457) |
| LE_13-9 | 1 l | Llanrwst WWTP | 2201 | RVA, RVC, PBV | NoVGII (1,251) |
| LI_11-10 | 1 l | Llanrwst WWTP | 859 | PBV | NoVGII (detected) |
| LE_11-10 | 1 l | Llanrwst WWTP | 5433 | NoVGI, RVA, RVC, PBV, AsV | NoVGII (50,180) |
| SW | 50 l | Morfa beach | 243 | - | - |
| Sed1 | 60 g | Morfa beach | 550<sup>d</sup> | - | - |
| Sed2 | 60 g | Morfa beach | 550<sup>d</sup> | - | - |

<sup>a</sup> LI: sewage influent; LE: sewage effluent; SW: estuarine surface water; Sed: estuarine sediment
<sup>b</sup> RVA: rotavirus A; RVB: rotavirus B; PBV: picobirnavirus; SaV: sapovirus; NoVGI: norovirus genogroup I; AsV: astrovirus
<sup>c</sup> Samples were tested with qRT-PCR for the following targets: NoVGI, NoVGII, SaV, HAV, HEV. Results reported in genome copies per liter (gc/l), NoVGII was detected below limit of quantification (approx. 200 gc/l) in sample LI\_11-10. Nov GII was the only target virus detected by qRT-PCR.
<sup>d</sup> Samples Sed1 and Sed2 were assembled together into the contig dataset Sed.

### Summary of viral diversity

The virus taxonomic diversity present in each sample was assessed by comparison of curated read and contig datasets with both the RefSeq Viral protein database and the non-redundant protein database of NCBI, using Diamond blastx (14) and lowest common ancestor taxon assignment with Megan 6 (15). For wastewater samples LI_13-9, LE_13-9 and LE_11-10, two libraries were processed (indicated with _1 and in the dataset names) and one each for the wastewater influent sample LI_11-10, the surface water sample (SW) and two sediment samples (Sed1 & Sed2). This section focuses on those reads and contigs that have been assigned to the viral fraction exclusively, disregarding sequences of cellular or unknown origin.

The wastewater samples showed a greater richness of known viruses and had a larger number of curated contigs than the surface water and sediment samples (Figures 2 & 3). At the viral family level, between 14 and 34 groups were observed for wastewater influent and effluent samples, including the unclassified levels, 12 for the surface estuarine water sample, and 11 and 5 for the sediment samples Sed1 and Sed2, respectively. The unclassified viruses and unassigned bins are indicated in red in Figure 2 and made up the majority of known reads in the estuarine sediment samples. In most of the viromes, dsDNA and ssDNA virus families were present, despite having performed a DNase treatment after viral nucleic acid extraction (Figures 2 &3). These families represented only a minor (<5%) proportion of the total assigned reads with a few exceptions. In wastewater influent sample LI_11-10, reads assigned to the dsDNA family *Papillomaviridae* accounted for 61% of the total and these reads were assembled into a single contig representing a near-complete betapapillomavirus genome. In the surface water sample reads assigned to the ssDNA families *Circoviridae* and *Microviridae* represented 50% and 12% of the total, respectively, assembling into contigs representing a significant proportion of the genome. The presence of both ssDNA and dsDNA virus signatures in all datasets is most likely due to incomplete digestion of the viral DNA with the DNase Max kit.

**Figure 2:**
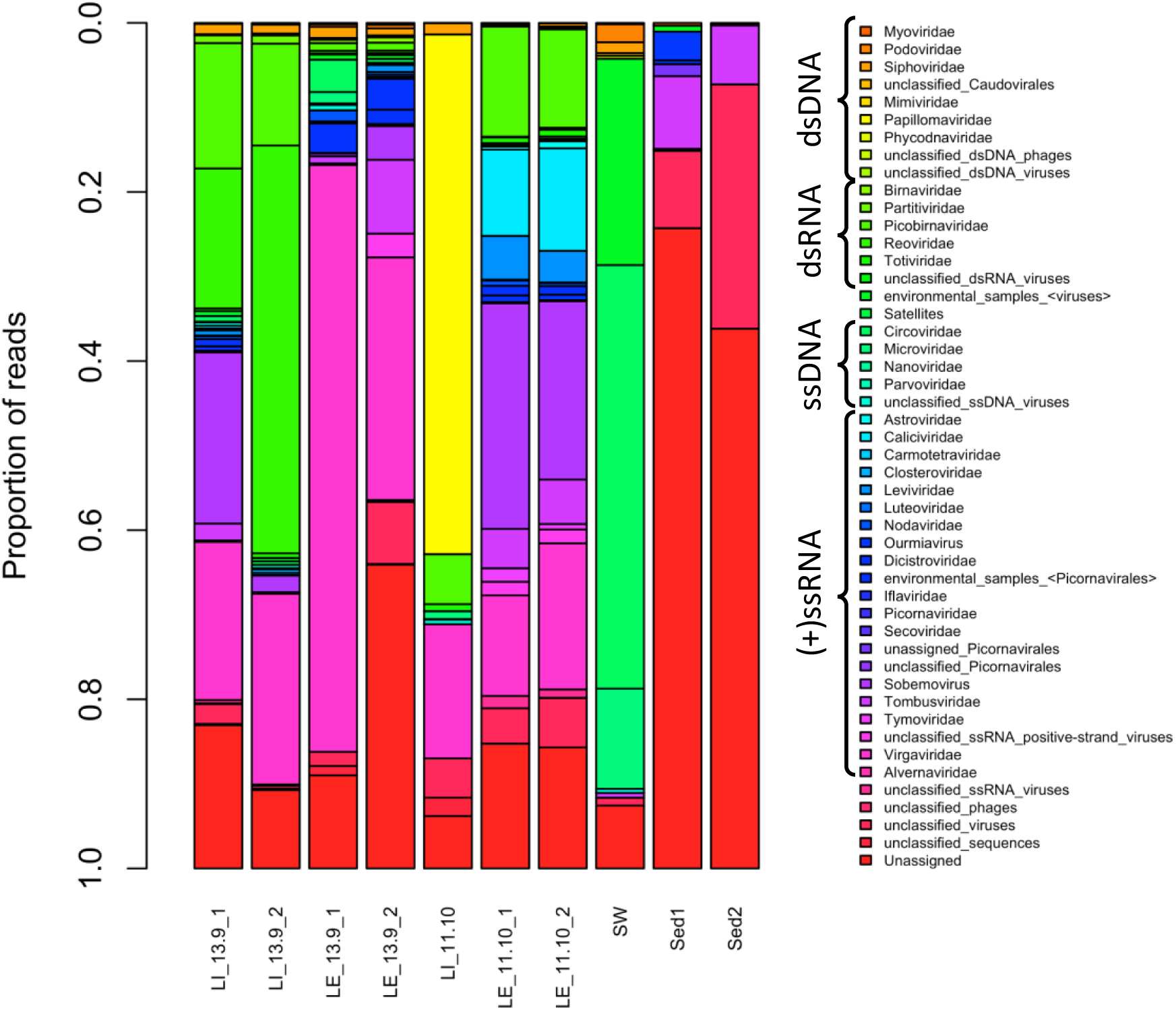
Taxonomic distribution of curated read data (relative abundance) at the virus family level. Reads were assigned to a family or equivalent group by Megan6 using a lowest common ancestor algorithm, based on blastx-based homology using the program Diamond with the RefSeq Viral protein database (version January 2017) and the non-redundant protein database (version May 2017). Only viral groupings are shown. LI: sewage influent; LE: sewage effluent; SW: estuarine surface water; Sed: estuarine sediment.

**Figure 3:**
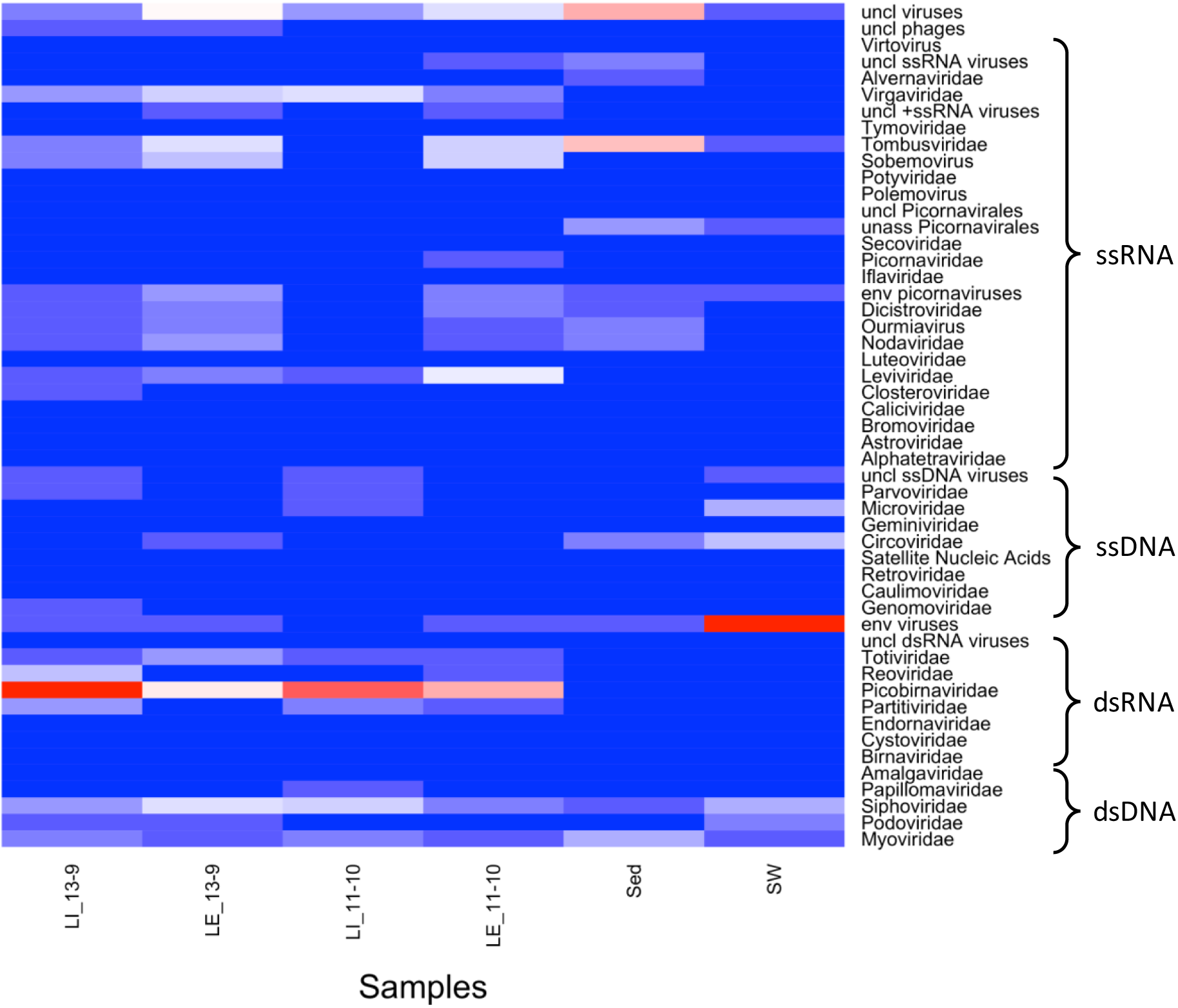
Heatmap of viral richness at the family level per sample. Heatmap colors range from blue (taxon not present or at low relative abundance in sample) over white to red (taxon present at high relative abundance in sample). Contigs larger than 300 nt were assigned to a family or grouping by Megan6 using a lowest common ancestor algorithm, based on blastx-based homology using the program Diamond with the RefSeq Viral protein database (version January 2017) and the non-redundant protein database (version May 2017). Only those families/groups comprising large contigs (>1000 nt) or with contigs mapping to viral signatures genes (e.g. capsid, RNA-dependent RNA polymerase) were retained. LI: sewage influent; LE: sewage effluent; SW: estuarine surface water; Sed: estuarine sediment.

The families of dsRNA viruses present in these datasets were *Totiviridae* (fungi and protist hosts), *Reoviridae* (invertebrate, vertebrate & plant hosts), *Picobirnaviridae* (mammals), *Partitiviridae* (fungi & protists) and *Birnaviridae* (vertebrates and invertebrates), with a small number of reads and contigs recognized as unclassified dsRNA viruses (Figures 2 & 3). None of these groups were present in all libraries, but totivirus and picobirnavirus signatures were present in all wastewater samples and reoviruses were found in three out of the four wastewater samples. *Partitiviridae* signatures were only found in the wastewater LE_11-10 and LI_13-9 samples, while *Birnaviridae* reads were only present in the wastewater LE_13-9 libraries. The sediment and surface water samples did not have detectable levels of dsRNA virus sequences.

Positive sense ssRNA viruses were the most diverse class of viruses present in these datasets. The family *Tombusviridae,* which groups plant viruses with monopartite or bipartite linear genomes (16), was present in all samples with the sole exception of the wastewater influent sample LI_11-10 (Figures 2 & 3). Virus signatures belonging to the family *Virgaviridae,* representing plant viruses, were present in all wastewater samples at comparable levels. Other highly represented families or groupings were the families *Dicistroviridae* (invertebrate hosts), *Nodaviridae* (invertebrate & vertebrate hosts) and the bacteriophage family *Leviviridae,* the plant virus genus *Sobemovirus,* and the groupings of “unclassified ssRNA positive-strand viruses” and several unclassified/unassigned/environmental members of the order *Picornavirales.* Sediment sample Sed1 was the only sample with signatures of the family *Alvernaviridae,* which has as its sole member the dinoflagellate virus Heterocapsa circularisquama RNA virus 01. The wastewater effluent sample LE_11-10 and influent sample LI_13-9_1 were the only samples with calicivirus signatures, and sample LE_11-10_1 and LE_1-10_2 were the only samples with *Astroviridae* reads (vertebrate host). Several families of the order *Picornavirales* were detected in the wastewater samples at different levels in different samples, and a small number of unassigned picornaviruses was detected in the surface water sample (SW).

We did not observe any known negative sense (-) ssRNA viruses in any of the sequencing libraries, but it is possible that some of the unaffiliated viral contigs belong to this class. The known human pathogenic (-) ssRNA viruses are enveloped (16) and predicted to degrade more rapidly than the non-enveloped enteric viruses, especially in wastewater (17, 18). We cannot rule out the possibility that (-) ssRNA viruses were present, but were removed by our sampling protocol.

The general wastewater viral diversity found here is similar to that reported previously. Those studies that investigated RNA viruses found both bacterial and eukaryotic viruses, with a high abundance of plant viruses of the family *Virgaviridae,* which includes the tobamovirus pepper mild mottle virus (11, 19). The families of viruses with potential human hosts found in previous metagenomics studies of sewage include *Astroviridae, Caliciviridae, Picobirnaviridae* and *Picornaviridae* (13, 19–21), of which only picobirnaviruses were recovered in all wastewater viromes in this study. In contrast, members of the family *Reoviridae,* represented by the genus *Rotavirus,* were found in three out of our four wastewater samples, but were not detected in many of the previous studies (19–21).

### Potential human pathogenic viruses

An important aim of this study was to investigate the presence and genomic diversity of potential human pathogenic RNA viruses in different sample types within the river catchment area. To minimize miss-assignments of short sequences to taxa, we used the assembled, curated contig dataset and looked for contigs representing near-complete viral genomes.

### Presence of a norovirus GI.2 genome

We were particularly interested in finding norovirus genomes in order to explore the genomic diversity of these important and potentially abundant pathogens originating from sewage and disseminated in watercourses, with implications for shellfisheries and recreational waters. This is of relevance due to known issues of sewage contamination in the region (22). Members of the genus *Norovirus* (family *Caliciviridae)* are non-enveloped, icosahedral (+)ssRNA viruses with a linear, unsegmented ~7.6 kb genome encoding three ORFs (16). These viruses are divided into different genogroups of which GI and GII are associated with human gastroenteritis (23, 24). Noroviruses are identified routinely by qRT-PCR, providing an opportunity here to examine correlations between qRT-PCR and metaviromic data.

We only found norovirus signatures in the libraries of wastewater effluent sample LE_11-10. These reads assembled into a single contig of 7,542 bases, representing a near-complete norovirus genome (GenBank accession number MG599789). Read mapping showed an uneven coverage over the genome length between 18x and 745x (13,165 reads of library 1 and 8986 reads of library 2). Based on this mapping, we performed variant calling and the consensus sequence was corrected in cases where the variant was present in more than 85% of the reads. To our knowledge, this is the only metagenome-derived, environment-associated (i.e. non-host associated) near-complete norovirus genome sequence deposited in a public database (INSDC nuccore database was searched for norovirus, txid142786 sequences > 5000 nt).

A BLASTN search revealed two close relatives to our wastewater-associated norovirus genome, norovirus Hu/GI.2/Jingzhou/2013401/CHN (KF306212) which is 7740 bases in length (25), displaying a nucleotide sequence identity of 99% over 99% of the genome length, and norovirus Hu/GI.2/Leuven/2003/BEL (FJ515294) at 95% sequence identity over 99% of the alignment length (Figure 4). From the 5’ end of our norovirus contig, 62 bases were missing compared with Hu/GI.2/Jingzhou/2013401/CHN and from the 3’ end 165 bases and the polyA tail were not present. We compared the sequence of our norovirus with Hu/GI.2/Jingzhou/2013401/CHN base by base and observed 81 SNPs and no other forms of variation. Of the SNPs, only eight were non-synonymous resulting in five different amino acids incorporated in the non-structural polyprotein (ORF1); one in the major capsid protein (ORF2) and two in the minor structural protein (ORF3). According to the current classification criteria, this level of similarity places our assembled genome in genogroup GI, genotype GI.2, with only a single amino acid different between the major capsid protein (MCP) of Hu/GI.2/Jingzhou/2013401/CHN and the genome assembled here.

**Figure 4:**
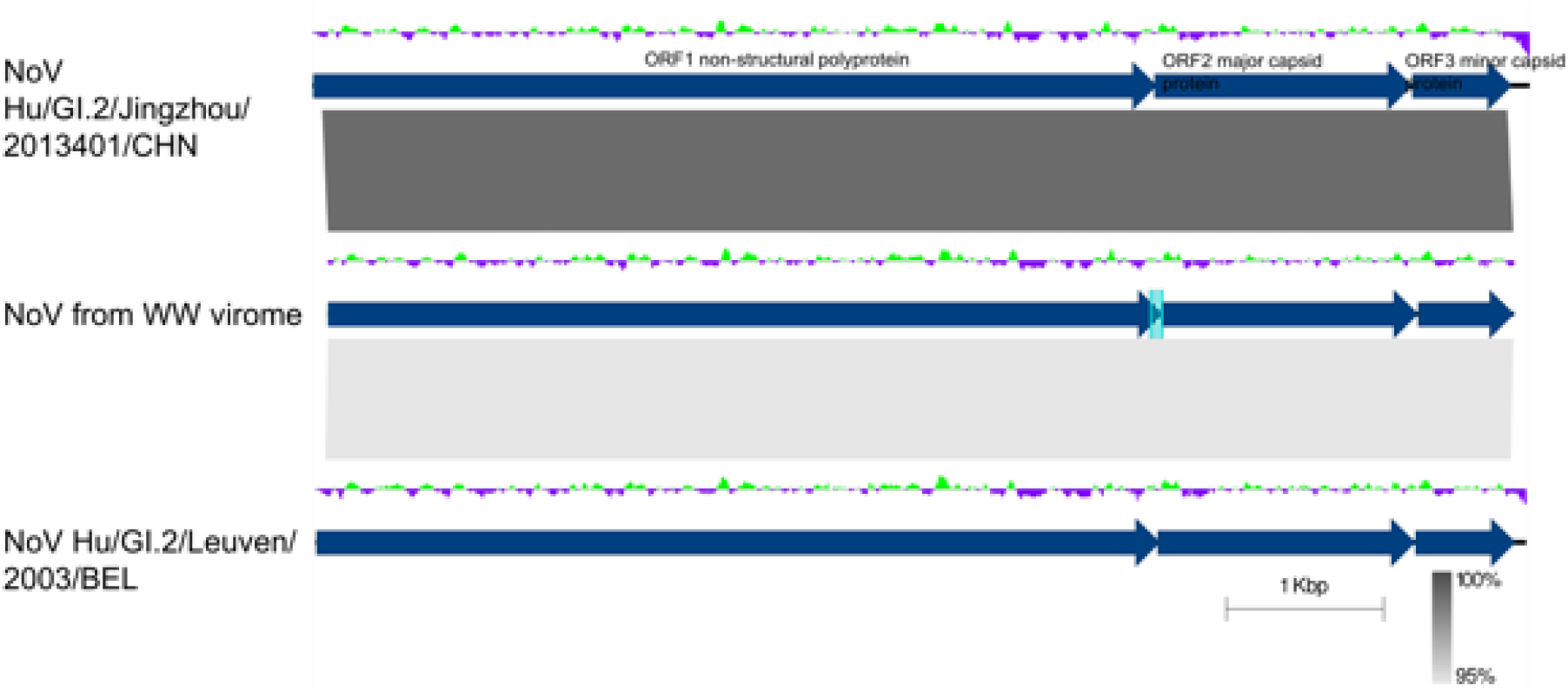
Pairwise genome comparison between the virome norovirus genome (middle) and its closest relatives, Norovirus Hu/GI.2/Jingzhou/2013401/CHN and Norovirus Hu/GI.2/Leuven/2003/BEL. BLASTN similarity is indicated in shades of grey. ORFs are delineated by dark blue arrows. The deviation from the average GC content is indicated above the genomes in a green and purple graph. The qRT-PCR primer binding sites for the wastewater-associated genome are indicated by light blue rectangles. The figure was created with Easyfig (81).

We tested the genotype grouping of our genome in a whole genome phylogeny with all complete genome sequences of genogroup I available in GenBank. The phylogenomic tree clearly delineated the different genotypes within genogroup GI, placing the newly-assembled genome within genotype GI.2, with the reference isolate for GII used as an outgroup (Figure 5).

**Figure 5:**
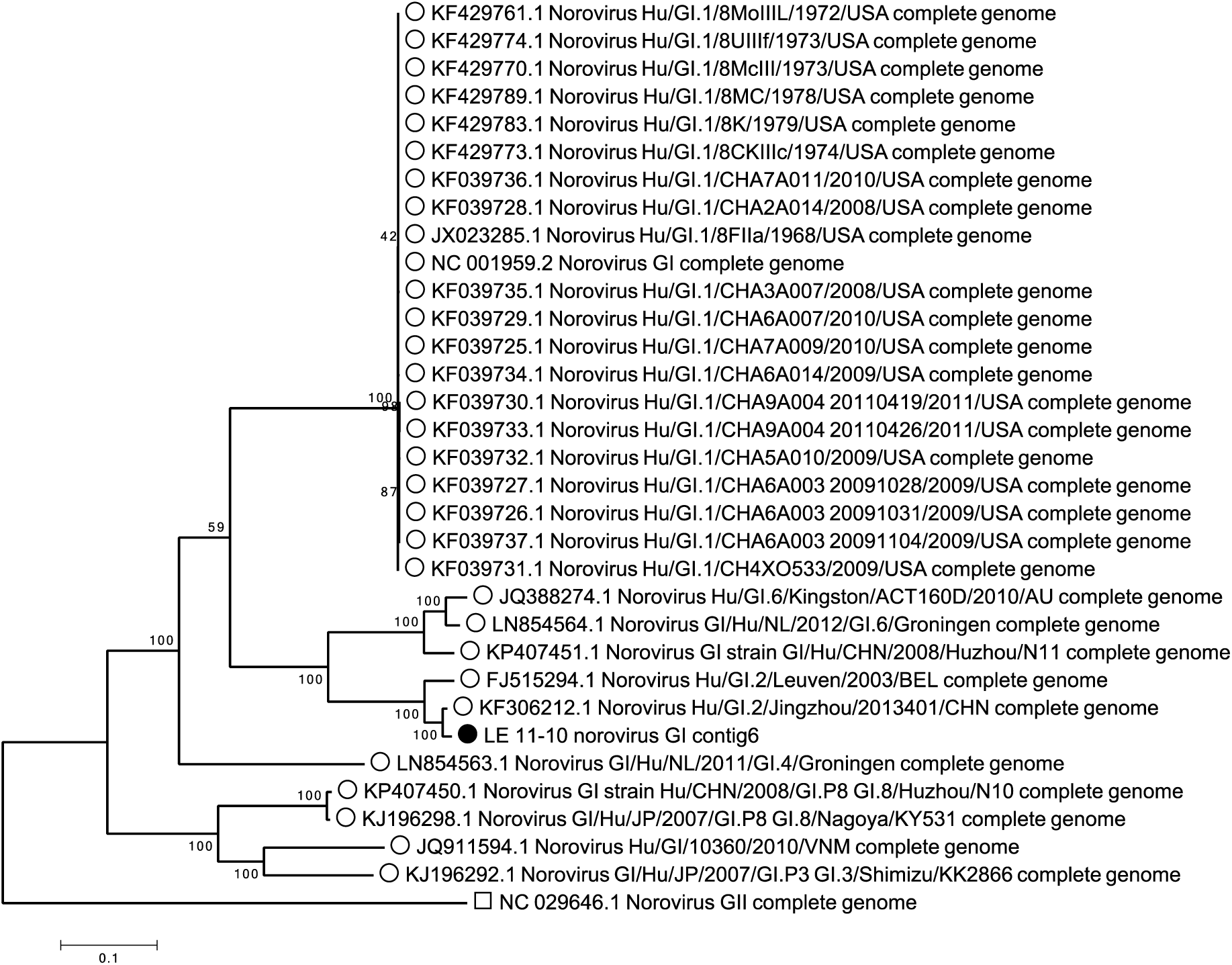
Maximum Likelihood phylogenetic tree of norovirus genomes belonging to genogroup GI, with the norovirus GII reference genome as an outlier. The nucleotide sequences were aligned with MUSCLE and the alignment was trimmed to the length of the virome sequence LE_11-10 contig 6, resulting in 7758 positions analysed for tree building. The Maximum Likelihood method was used with a Tamura Nei model for nucleic acid substitution. The percentage of trees in which the associated taxa clustered together is shown next to the branches. The scale bar represents the number of substitutions per site.

For further validation, the full genome of the novel norovirus GI was recovered using RT-PCR. However, the amplicon could not be ligated into a plasmid and hence was not fully sequenced.

### Presence of diverse rotavirus segments in wastewater samples

Rotaviruses are segmented dsRNA viruses belonging to the family *Reoviridae,* causing gastroenteric illness in vertebrates and are transmitted through the faecal-oral route (16). Read signatures assigned to the genus *Rotavirus* were found in three of the four wastewater samples (all but LI_11-10). Wastewater influent sample LI_13-9 contained the most signatures with approximately 75,000 reads, assembled into 120 contigs, representing genome fragments of 10 out of the 11 rotavirus segments. At the species level, these genome fragments were assigned to either the species *Rotavirus A* or *Rotavirus C.* Comparing the amino acid sequences of the predicted proteins, some contigs showed high levels of identity (>88%) with either the segments of rotavirus A (RVA) or rotavirus C (RVC) reference genomes as available in the RefSeq database (26, 27), while others showed a lower identity with a variety of RVC isolates only. The segmented genome nature and the possibility of segment exchange make it difficult to confidently identify the number of rotavirus types present in this sample. Given the amino acid similarities with both RVA and RVC types (Supplementary Table 1), we suggest there are at least two, and possibly three types present here.

Using the RotaC 2.0 typing tool for RVA, and blast-based similarity to known genotypes, we have typed the rotavirus genome segments found here (Table 2). The combined genomic make-up of the RV community in sample LI_13-9 was G8/G10/Gx-P[1]/P[14]/P[41]/P[x]-I2/Ix-R2/Rx-C2/Cx-M2/Mx-A3/A11/Ax-Nx-T6/Tx-E2/Ex (28, 29). The potential hosts for each segment were derived from the hosts of the closest relatives. This analysis showed that the RVA viruses were possibly infecting humans (through zoonotic transmission) or cattle, while the RVC viruses were most likely porcine (Table 2). However, due to the genomic diversity of the segments found here, particularly for RVC genome fragments, we cannot rule out alternative hosts.

**Table 2:** Rotavirus A and C genome information and its detection in the LI_13-9 sample dataset.

| Genome segment | Length (nt) | Protein | Predicted function | # contigs | Putative genotypes | Potential hosts<sup>a</sup> |
| --- | --- | --- | --- | --- | --- | --- |
| **RVA** |  |  |  |  |  |  |
| Segment 1 | 3302 | VP1 | RNA-dependent RNA polymerase | 7 | R2 | Human, cow |
| Segment 2 | 2693 | VP2 | core capsid protein | 1 | C2 | Human |
| Segment 3 | 2591 | VP3 | RNA capping protein | 1 | M2 | Human, sheep |
| Segment 4 | 2363 | VP4 | outer capsid spike protein | 3 | P[1], P[41], P[14] | Human, pig, alpaca, monkey |
| Segment 5 | 1614 | NSP1 | interferon antagonist protein | 6 | A3, A11 | Human, cow, pig, deer |
| Segment 6 | 1356 | VP6 | inner capsid protein | 1 | I2 | Human |
| Segment 7 | 1105 | NSP3 | translation effector protein | 4 | T6 | Human, dog, cow |
| Segment 8 | 1059 | NSP2 | viroplasm RNA binding protein | 0 | - | - |
| Segment 9 | 1062 | VP7 | outer capsid glycoprotein | 2 | G10, G8 | Cow, Human |
| Segment 10 | 751 | NSP4 | enterotoxin | 1 | E2 | Human, cow |
| Segment 11 | 667 | NSP5;6 | phosphoprotein; non-structural protein | 0 | - | - |
| **RVC** |  |  |  | (contigs RVCX) |  |  |
| Segment 1 | 3309 | VP1 | RNA-dependent RNA polymerase | 7 (0) | Rx | Pig, cow |
| Segment 2 | 2736 | VP2 | core capsid protein | 4(2) | Cx | Pig, dog |
| Segment 3 | 2283 | VP4 | outer capsid protein | 2 (4) | P[x] | Pig |
| Segment 4 | 2166 | VP3 | guanylyltransferase | 6 (0) | Mx | Pig |
| Segment 5 | 1353 | VP6 | inner capsid protein | 1 (0) | Ix | Pig |
| Segment 6 | 1350 | NSP3 |  | 0 (1) | Tx | Human |
| Segment 7 | 1270 | NSP1 |  | 0 (2) | Ax | Pig, dog |
| Segment 8 | 1063 | VP7 | outer capsid glycoprotein | 0 (2) | Gx | Pig |
| Segment 9 | 1037 | NSP2 |  | 2 (0) | Nx | Pig |
| Segment 10 | 730 | NSP5 |  | 0 (0) | - | - |
| Segment 11 | 613 | NSP4 | enterotoxin | 0 (4) | Ex | Pig |

<sup>a</sup> Potential hosts are defined as the hosts of the reference rotavirus sequence with the highest similarity to the contigs found in the virome sample LI\_13-9.

### Partial genomes of other potentially pathogenic RNA viruses

In sample LI_13-9, a small contig of 347 bases was found that was 94% identical at the nucleotide level to the Sapovirus Mc2 ORF1 (AY237419), in the family *Caliciviridae.* We have also identified four contigs of approximately 500 bases in sample LE_11-10 that resembled most closely the Astrovirus MLB2 isolates MLB2/human/Geneva/2014 (KT224358) and MLB2-LIHT (KX022687) at 99% nucleotide identity. In addition, we identified several reads and contigs assigned to the family *Picornaviridae* which comprises a diverse set of enteric viruses, but the closest relatives in the databases were metagenomically assembled or unidentified picornaviruses.

### Picobirnaviruses showed a high prevalence in wastewater

All the wastewater virome libraries contained signatures assigned to the dsRNA family *Picobirnaviridae,* genus *Picobirnavirus* (Figure 2) and these reads assembled into between 42 (LE_13-9) and 510 (LI_13-9) contigs. Both picobirnavirus genome segments, segment 1 containing two hypothetical proteins and segment 2 on which the RNA-dependent RNA polymerase (RdRP) is encoded, were observed in the samples. The contigs showed little sequence similarity with the reference genome *Human picobirnavirus* (RefSeq segment accession numbers NC_007026.1 and NC_007027.1). Phylogenetic analysis of a partial region of the predicted RdRPs in the virome contigs was not able to resolve any cluster or evolutionary origin (Figure 6A). Picobirnavirus RdRPs from human, animal and environmental isolates, as well as the majority of the virome sequences were grouped in one large, unsupported cluster that showed relatively little genomic diversity. While many picobirnaviruses have been isolated from humans with gastroenteritis, a review of the known cases suggested that picobirnaviruses are probably not the main cause of acute diarrhoea and are secondary pathogens with potential synergistic effects (30). A qRT-PCR-based investigation into the suitability of human picobirnaviruses as indicators of human faecal contamination, showed that they were not present in a sufficient proportions of tested samples to be good water quality indicators (31), but their high diversity in our sample set warrants further investigation for their use as water quality markers using metaviromic methods.

**Figure 6:**
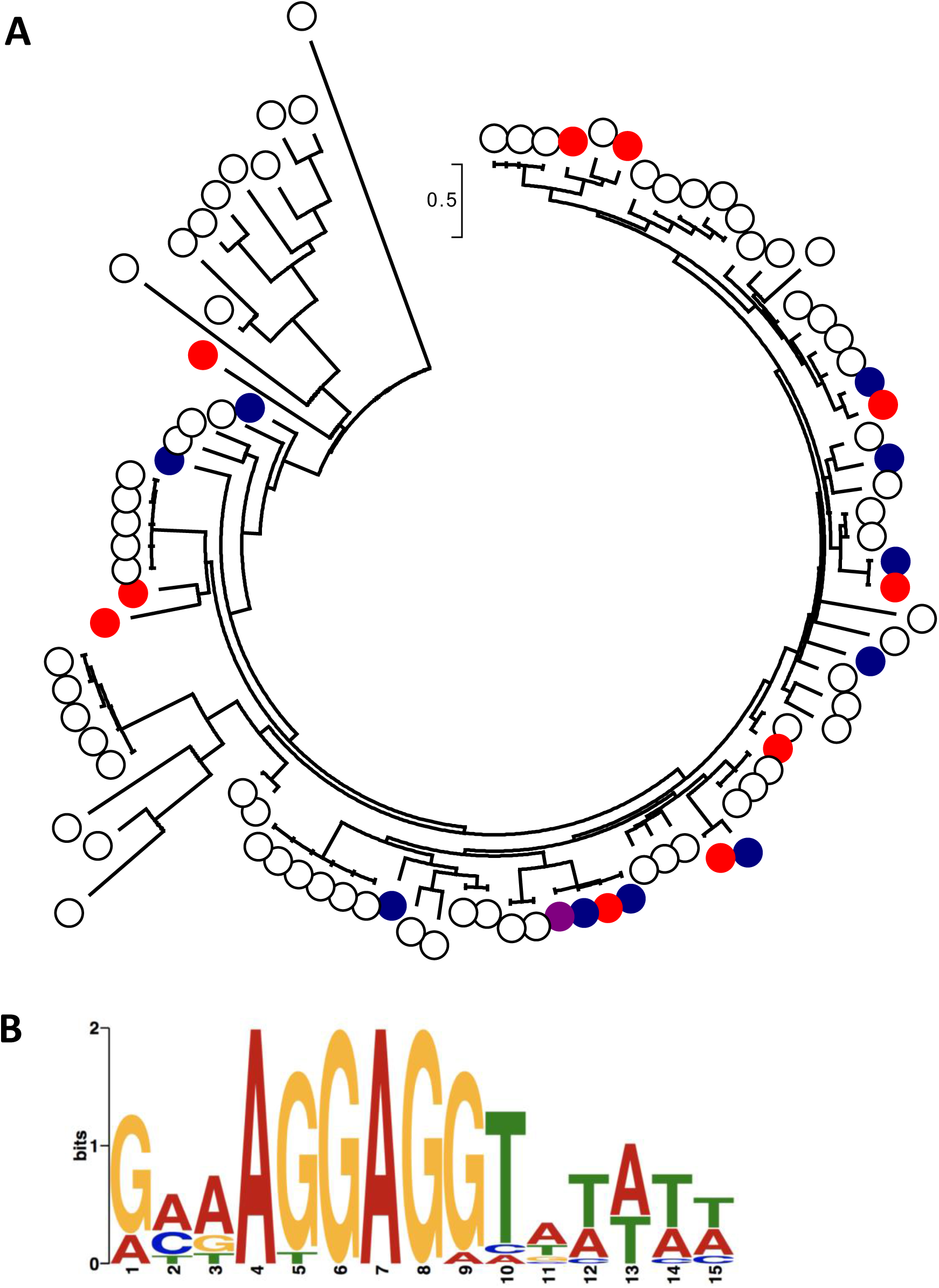
Picobirnavirus diversity. A) Maximum likelihood phylogenetic tree of RdRP amino acid sequences of isolated and virome picobirnaviruses. Sequences from isolates are indicated with white dots, virome-derived sequences with filled-in coloured dots, sample LI_11-10 in purple, sample LE_11-10 in blue, and sample LI_13-9 in red. Sequences were aligned using MUSCLE providing 114 amino acid positions for tree generation. The Maximum Likelihood was used with a JTT matrix-based model. The scale bar represents the number of substitutions per site. Bootstrap values of all branches were low. B) Predicted ribosome binding site consensus sequence from extracted 5’ UTRs, logo produced by the MEME-suite.

A recent study of picobirnaviruses produced the hypothesis that these viruses do not infect mammals, but are a new family of RNA bacteriophages, based on the presence of bacterial ribosome binding sites (RBS) upstream of the coding sequences (CDS) (32). To test this hypothesis, we extracted all contigs with amino acid similarity to the RdRP or capsid protein of known picobirnaviruses, annotated the CDS and extracted the upstream 21 nucleotides from the transcription start site. In the 233 contigs found, 71 partial CDSs were predicted from which we extracted 17 5’ UTRs (untranslated regions), discarding those partially annotated CDSs missing the transcription start site. We discovered the 6-mer motif AGGAGG (Figure 6B) in 100% of the upstream sequences, similar to the frequency reported by Krishnamurthy and Wang (32), who found at least a 4-mer RBS in 100% of the 98 picobirnavirus 5’ UTRs investigated. In contrast, the different families of eukaryotic viruses analysed in that study only showed a low incidence of RBSs, which were mostly 4-mers. Our findings, therefore, support the hypothesis that picobirnaviruses are bacteriophages and we suggest that they belong to a novel RNA bacteriophage family with a high level of genomic diversity.

## Discussion

We set out to explore the possibility of using viromics to find human pathogenic RNA viruses in the environment. We have been successful in identifying several potentially human pathogenic, including potentially zoonotic, viral genomes from the wastewater samples, but did not find any in the surface estuarine water and sediment samples. The absence of signatures does not necessarily mean that there are no pathogenic viruses present in water or sediment, but only that their levels could be below our limit of quantification for qPCR (approximately 200 gc/l).

It is important to note here that during the RNA extraction process, many biases could have been introduced leading to a lower recovery of input viruses. Samples were first concentrated from volumes of 1 l (wastewater) or 50 l (surface water) down to 50 ml using tangential flow filtration (TFF) at a molecular weight cut-off of 100 kDa, followed by PEG 6000 precipitation. These samples were diluted in fresh buffer, filtered through syringe filters of 0.22 μm pore size and then treated with nuclease to remove free DNA and RNA. Previous research has shown that while any enrichment method aimed at fractionating the viral and cellular components will decrease the total quantity of viruses, a combination of centrifugation, filtration and nuclease treatment increases the proportion of viral reads in sequencing datasets (33). After implementing these steps, we used the MO BIO PowerViral® Environmental DNA/RNA extraction kit for viral RNA extraction, which has previously been shown to perform best overall in spiking experiments with murine norovirus, in terms of extraction efficiency and removal of inhibitors (34). The kit has, however, given low recoveries of viruses from sediment (35).

We did not perform an amplification step before library construction with the NEBNext Ultra Directional RNA Library Prep Kit for Illumina, to retain the genome sense and strand information. Instead, we increased the number of cycles of random PCR during library preparation from 12 to 15 to counteract the low input quantity of RNA (< 1 ng). The random amplification during library construction led to a trade-off in which genome strand information was gained for a loss of quantitative power, making it difficult to compare abundances of viral types within and across libraries. This random PCR-based bias has been highlighted before, but the proposed solution of using library preparation protocols which limit the use of PCR are only feasible with high amounts of input nucleic acid (36), which we have not found to be possible when processing environmental/wastewater samples to generate RNA metaviromes.

A critical issue to highlight here, is the inclusion of controls in our sequencing libraries in order to identify potential contaminants and their origins, as has been suggested previously (37, 38). There have been multiple reports of false positive genome discoveries, in particular the novel parvovirus-like hybrid in hepatitis patients that was later revealed to originate from the silica-based nucleic acid extraction columns (39–41). In this study, we included a positive control that comprised bacterial cells *(Salmonella enterica* serovar Typhimurium isolate D23580 RefSeq accession number NC_016854) and mengovirus (36), an RNA virus that serves as a process control, as well as two negative controls, an extraction control and a library preparation control. Analysis of the control libraries showed that while the *Salmonella* cells and DNA were successfully removed from the positive control sample by the enrichment protocol, the mengovirus was not recovered. Subsequent qRT-PCR analysis revealed that the mengovirus remained detectable in the pre-processing stages of the extraction, but was lost after RNase treatment (data not shown). Inclusion of an inactivation step of the DNase at 75°C potentially exacerbated the effect of the RNase step. Consequently, it is likely that we have missed viral types during the extraction process despite having still managed to recover an RNA metavirome harbouring substantial diversity.

Further examination of the HiSeq and MiSeq control datasets revealed a wide range of contaminant signatures of prokaryotic, eukaryotic and viral origin, making up 45M read pairs per control on the HiSeq platform and 1M read pairs for the MiSeq, even though the 16S and 18S rRNA PCR and RT-PCR reactions showed no visible bands on an agarose gel. Most bacterial contaminant reads belonged to the phyla *Proteobacteria, Actinobacteria* and *Firmicutes.* The most abundant genera included *Corynebacterium, Propionibacterium, Sphingomonas, Ralstonia, Pseudomonas, Streptomyces, Staphylococcus* and *Streptococcus* which have in the past been identified as common lab contaminants (42). Within the eukaryotic signatures, human-derived reads, *Beta vulgaris* and *Anopheles* reads were the most prevalent, pointing towards potential cross-contamination of the sequencing libraries. A small number of virus signatures were also identified, with the most prominent being *Feline calicivirus* and *Dengue virus.* The presence of the calicivirus was traced back to the library preparation kits after the libraries were reconstructed and resequenced. The dengue virus signature was a <100 nt sequence which was co-extracted in all the samples and potentially originated in one of the reagents or spin extraction column. All sequences present in the controls were carefully removed from the sample datasets during the quality control stage of the bioinformatics processing before further analysis. For future experiments, we will omit the RNase treatment step during extraction and filter out any contaminating ribosomal RNA or cellular-derived mRNA sequences as part of the bioinformatic quality control workflow.

Our results show that while contamination is an issue when dealing with low biomass samples, the combination of increased random PCR cycles during library preparation, deep sequencing (i.e. HiSeq rather than MiSeq) and computational subtraction of control sequences provides data of sufficient quantity and quality to assemble near-complete RNA virus genomes *de novo.*

### Norovirus

Noroviruses are one of the most common causes of gastrointestinal disease in the developed world, with an incidence in the UK estimated as approaching 4 million cases per annum (43). The genotype most commonly associated with disease is GII.4 (44–46) which was not detected in the metaviromes generated here.

We retrieved one norovirus GI genome, assembled from 22,151 reads, in wastewater effluent sample LE_11-10. This finding was in direct conflict with the qRT-PCR analysis of this sample which did not detect any NoV GI signatures (Table 1). In contrast, NoV GII signatures were detected by qRT-PCR, but no NoV GII genomes or genome fragments were observed in the virome libraries. One hypothesis to explain the discrepancy between PCR and viromics approaches lies in the differences in extraction protocol. For qRT-PCR, no viral enrichment step was performed and RNA was not extracted with the PowerViral kit. Therefore, NoV GII could have been lost before virome sequencing, as was the process control mengovirus. An alternative hypothesis is that the NoV GII signatures detected during qRT-PCR were derived from fragmented RNA or from particles with a compromised capsid. In both these cases, the RNA would not be detected in the virome data because of the RNase preprocessing steps implemented in the enrichment/extraction protocol. This calls into question the reliance of qRT-PCR for NoV detection and whether the detected viruses are infectious or merely remnants of previous infections. Further research using, for example, capsid integrity assays combined with infectious particle counts will need to be conducted to assess the validity of qRT-PCR protocols for norovirus detection.

The inability to identify NoV GI with qRT-PCR might be related to the mismatched base present in the forward primer sequence used for detection. We subsequently conducted a normal, long-range PCR to validate the detection of this genotype, and this yielded a fragment of the correct size, but we were unable to clone and sequence this fragment. While the known NoV GI.2 genotypes do not have a mismatch in the qRT-PCR probe sequence, it is possible that the genome recovered in this study fell below the limit of detection using the ISO standard primer/probe combination (ISO/TS 15216-2:2013). In a recent study, researchers designed an improved probe and observed lower Ct values and a lower limit of detection for GI.2 strains from waterborne samples (47). Viromics as a means of investigating water samples for the presence of norovirus, does have the advantage of demonstrating the presence of an undegraded genome, provided the sample processing requirements do not lead to excessive loss of virus particles resulting in false negatives. Certainly, time and cost permitting, viromics is a useful adjunct to qPCR for samples that are deemed particularly important or critical for determination of intact viral genome presence.

Due to the virtual impossibility of culturing noroviruses in the lab, many studies have used male-specific coliphages such as MS2 and GA, which are ssRNA phages belonging to the family *Leviviridae,* as alternative model systems (48, 49). Interestingly, while some levivirus signatures were present in all wastewater samples (< 500 reads), we observed a striking co-occurrence of these viruses with norovirus signatures in both libraries of sample LE_11-10 (> 2500 reads). The most commonly observed viruses in this sample were *Pseudomonas* phage PRR1, an unclassified levivirus, and *Escherichia* phages FI and M11 in the genus *Allolevivirus.* Further studies with more samples and replicates will indicate whether there is a significant correlation between the presence of leviviruses and noroviruses in water samples. Furthermore, the higher abundance of alloleviviruses compared with MS2-like viruses could indicate that the former might be more relevant as model systems for noroviruses.

### Rotavirus

Rotaviruses are, like noroviruses, agents of gastroenteritis, but the disease is commonly associated with children under the age of 5 where severe diarrhoea and vomiting can lead to over 10,000 hospitalizations per year in England and Wales (50). Since the introduction of the live-attenuated vaccine Rotarix, the incidence of gastroenteritis in England has declined, specifically for children aged <2 and during peak rotavirus seasons (51–53). Therefore, the discovery of a diverse assemblage of rotavirus genome segments in the wastewater samples here was less expected than the norovirus discovery. While we were unable to recover the genome of the vaccine strain, our genomic evidence suggests that at least one RVA and one RVC population were circulating in the Llanrwst region in September 2016.

The genome constellation for the RVA segments in sample LI_13-9, G8/G10-P[1]/P[14]/P[41]-I2-R2-C2-M2-A3/A11-(N)-T6-E2-(H), is distinctly bovine in origin (28) (N and H segments not recovered in this study). The closest genome segment relatives based on nucleic acid similarity, however, have been isolated from humans (Table 2), possibly pointing towards a bovine-human zoonotic transmission of this virus (54). The same genomic constellation has been found recently when unusual G8P[14] RVA isolates were recovered from human strain collections in Hungary (55) and Guatemala (56), and isolated from children in Slovenia (57) and Italy (58). Cook and colleagues calculated that there would be approximately 5000 zoonotic human infections per year in the UK from livestock transmission, but many would be asymptomatic (59).

The origins of the RVC genome segments are more difficult to trace, because of lower similarity scores with known RVC isolates. The majority of the segments were similar to porcine RVC genomes, while others showed no nucleotide similarity at all, only amino acid similarity. An explanation for the presence of pig-derived rotavirus signatures could be farm run-off. While farm waste is not supposed to end up in the sewage treatment plant, it is likely that the RVC segments originate directly from pigs, not through zoonotic transfer. Run-off from fields onto public roads, broken farm sewer pipes or polluted small streams might lead to porcine viruses entering the human sewerage network, but we cannot provide formal proof from the data available. Based on the evidence, we hypothesize that there is one, possibly two, divergent strains of RVC circulating in the pig farms in the Llanrwst area.

## Conclusion

In this study, we investigated the use of metagenomics for the discovery of RNA viruses circulating in watercourses. We have found RNA viruses in all samples tested, but potential human pathogenic viruses were only identified in wastewater. The recovery of plant viruses in most samples points towards potential applications in crop protection, for example the use of metaviromics in phytopathogen diagnostics. However, technical limitations, including the amount of input material necessary and contamination of essential laboratory consumables and reagents, are currently the main bottleneck for the adoption of fine scale metagenomics in routine monitoring and diagnostics. The discovery of a norovirus GI and a diverse set of rotavirus segments in the corresponding metaviromes indicates that qPCR-based approaches can miss a significant portion of relevant pathogenic RNA viruses present in water samples. Therefore, metagenomics can, at this time, best be used for exploration, to design new diagnostic markers/primers targeting novel genotypes and to inform diagnostic surveys on the inclusion of specific additional target viruses.

## Materials & Methods

### Sample collection and processing

Wastewater samples were collected as part of a viral surveillance study described elsewhere (Farkas et al, in submission). Wastewater influent and effluent, 1l each, was collected at the Llanrwst wastewater treatment plant by Welsh Water (Wales, UK, Figure 1) on 12^th^ September (processed on 13-9, sample designations LI_13-9 and LE_13-9) and 10^th^ October 2016 (processed on 11-10, sample designations LI_11-10 and LE_11-10). The wastewater treatment plant uses filter beds for secondary treatment and serves approx. 4000 inhabitants. The estuarine surface water (50 L) sample (SW) was collected at Morfa Beach (Conwy, Wales, Figure 1) approx. 22 km downstream of the Llanrwst wastewater treatment plant on 19^th^ October and 2^nd^ of November 2016 at low tide (only the sample from November was used for sequencing as the October sample extract failed quality control). Together with the surface water sample, 90 g of the top 1-2 cm layer of the sediment was also collected (sample designations Sed1 for the October sample and Sed2 for the November sample).

The wastewater and surface water samples were processed using a two-step concentration method as described elsewhere (Farkas et al, in submission). In brief, the 1l (wastewater) and 50l (surface water) samples were first concentrated down to 50 ml using a KrosFlo^®^ Research IIi Tangential Flow Filtration System (Spectrumlabs, USA) with a 100 PEWS membrane. Particulate matter was then eluted from solid matter in the concentrates using beef extract buffer and then viruses were precipitated using polyethylene glycol (PEG) 6000. The viruses from the sediment samples were eluted and concentrated using beef extract elution and PEG precipitation as described elsewhere (35). The precipitates were eluted in 2-10 mL phosphate saline buffer, (PBS, pH 7.4) and stored at −80°C.

### Detection and quantification of enteric viruses with qRT-PCR

Total nucleic acids were extracted from a 0.5 mL aliquot of the concentrates using the MiniMag NucliSENS® MiniMag® Nucleic Acid Purification System (bioMérieux SA, France). The final volume of the nucleic acid solution was 0.05 mL (surface water and sediment) and 0.1 mL (wastewater samples). Norovirus GI and GII, sapovirus GI, and hepatitis A and E viruses were targeted in qRT-PCR assays as described elsewhere (60).

### Viral RNA extraction for metaviromic sequencing

Viral particles were extracted from the concentrated samples by filtration. In a first step, the samples were diluted in 10 ml of sterile 0.5 M NaCl buffer and incubated at room temperature (20°C) with gentle shaking for 30 min to disaggregate particles. The suspension was then filtered through a sterile, 0.22 μm pore size syringe filter (Millex, PES membrane). The sample was desalted by centrifugation (3200 x g, between 1 and 6h for different samples) in a sterilized spin filter (Vivaspin 20, 100 kDa molecular weight cut-off) and replacement of the buffer solution with 5 ml of a Tris-based buffer (10 mM TrisHCl, 10 mM MgSO4, 150 mM NaCl, pH 7.5). The buffer exchange was performed twice and the volume retained after the final spin was < 500 μl. The samples were then treated with Turbo DNase (20 Units; Ambion) and incubated for 30 minutes at 37°C, followed by inactivation at 75°C for 10 minutes. In a next step, all samples were treated with 80 μg RNase A (Thermo Fisher Scientific) and incubated at 37°C for 30 minutes. The RNase was inactivated with RiboLock RNase Inhibitor (Thermo Fisher Scientific) and the inactivated complex was removed by spin filtration (Vivaspin 500, 100 kDa molecular weight cut-off) and the samples centrifuged until the volume was approximately 200 μl. Viral DNA and RNA were co-extracted using the PowerViral Environmental DNA/RNA kit (MOBIO Laboratories) according to the manufacturer’s instructions. In this protocol, buffer PV1 was supplemented with 20 μl/ml betamercaptoethanol to further reduce RNase activity. The nucleic acid was eluted in 100 μl RNase-free water. The extracted viral DNA was degraded using the DNase Max kit (MOBIO Laboratories) according to the manufacturer’s instructions. The remaining viral RNA was further purified and concentrated by ethanol precipitation using 2.5 x sample volume of 100% ethanol and 1/10 volume of DEPC-treated Na-acetate (3 M). The quantity and quality of RNA was determined with Bioanalyzer Pico RNA 6000 capillary electrophoresis (Agilent Technologies). A positive and negative extraction control sample were processed alongside the main samples. The positive control samples contained *Salmonella enterica* serovar Typhimurium strain D23580 which is not found in the UK (61) and a process control virus mengovirus (60, 62).

The viral RNA extracts were tested for bacterial and eukaryotic cellular contamination using 16S and 18S rRNA gene PCR and RT-PCR, with primers e9F (63) and 519R (64), and primers 1389F and 1510R (65), for the 16S and 18S rRNA gene, respectively. Complimentary DNA was created using the SuperScript III Reverse Transcriptase (Invitrogen) with random hexamer primers according to the manufacturer’s instructions. (RT)-PCR was performed with the MyTaq Red Mix (Bioline) for 35 cycles (95°C for 45 sec, 50°C for 30 sec, 72°C 1 min 40 sec) and visualized on a 1% agarose gel. Samples were considered suitable for sequencing if no DNA bands were visible on the gel.

### Library preparation and sequencing

The library preparation and sequencing were performed at the University of Liverpool Centre for Genomics Research (CGR). Twelve dual indexed, strand-specific libraries were created using the NEBNext Ultra Directional RNA Library Prep Kit for Illumina, according to the manufacturer’s instructions. These libraries were pooled and sequenced at 2 × 150 bp read lengths on the Illumina HiSeq 4000 platform. This generated between 10 and 110 million paired reads per sample.

To confirm our results, a second set of libraries was constructed from new kits and a milliQ water samples was included as a library prep control. The thirteen resulting libraries were sequenced on the Illumina MiSeq platform at CGR, at 2 × 150 bp read lengths. These data were used for verification and control purposes only as sequencing depth was insufficient for the bioinformatics analyses described in the rest of the study.

### Bioinformatics

All command line programs for data analysis were run on the bioinformatics cluster of CGR (University of Liverpool) in a Debian 5 or 7 environment.

Raw fastq files were trimmed to remove Illumina adapters using Cutadapt version 1.2.1 using option -O 3 (66) and Sickle version 1.200 with a minimum quality score of 20 (67). Further quality control was performed with Prinseq-lite (68) with the following parameters: minimum read length 35, GC percentage between 5-95%, minimum mean quality 25, dereplication (removal of identical reads, leaving 1 copy), removal of tails of minimum 5 polyN sequences from 3’ and 5’ ends of reads.

The positive and negative control libraries described earlier were used for contaminant removal. The reads of the control samples were analysed using Diamond blastx (14) against the non-redundant protein database of NCBI (nr version November 2015). The blast results were visualised using Megan6 Community Edition (15). An extra contaminant file was created with complete genomes of species present at over 1000 reads in the positive and negative control samples. Then, bowtie2 (69) was used for each sample to subtract the reads that mapped to the positive control, negative control or contaminant file. The unmapped reads were used for assembly with SPAdes version 3.9.0 with kmer values 21, 31, 41, 51, 61, 71, and the options ‐‐careful and a minimum coverage of 5 reads per contig (70). The contig files of each sample were compared with the contigs of the controls (assembled using the same parameters) using blastn of the BLAST+ suite (71). Contigs that showed significant similarity with control contigs were manually removed, creating a curated contig dataset. The unmapped read datasets were then mapped against this curated contig dataset with bowtie2 and only the reads that mapped were retained, resulting in a curated read dataset.

The curated contig and read datasets were compared to the Viral RefSeq (release January 2017) and non-redundant protein (nr, release May 2017) reference databases using Diamond blastx at an e value of 1e-5 for significant hits (14, 72, 73). Taxon assignments were made with Megan6 Community Edition according to the lowest common ancestor algorithm at default settings (15). We have chosen the family level taxon assignments to represent the overall viral diversity, because there is generally little amino acid identity between viral families. The taxon abundance data were extracted from Megan6 and imported into RStudio for visualization (74). Genes were predicted on the assembled contigs with Prokka (75) using the settings - -kingdom Viruses and an e value of 1e-5. Multiple alignments of genes and genomes were made in MEGA7 using the MUSCLE algorithm at default settings (76, 77). The alignments were manually trimmed and phylogenetic trees were built using the Maximum Likelihood method in MEGA7 at the default settings. Upstream sequences of potential CDSs of prokka annotated picobirnaviruses were extracted using extractUpStreamDNA (https://github.com/ajvilleg/extractUpStreamDNA) and all 5’ UTRs and transcription start sites were manually verified in UGene (78). These extracted sequences were then subjected to a motif search using the MEME Suite (79, 80).

### Accession numbers

Read and contig datasets are available from NCBI under the following BioProject accession numbers, PRNJA421889 (wastewater data), PRNJA421892 (sediment data) and PRJNA421894 (estuarine water data). The NoV GI genome isolate was deposited in GenBank under accession number MG599789.

## Author contributions

EMA, KF, DJ, HA and AJM designed the experiments, EMA, KF, CH, performed the experiments, EMA analysed the data, EMA and KF wrote the manuscript and EMA prepared the manuscript for submission. All authors critically reviewed and edited the manuscript.

## Acknowledgements

This study was funded by the Natural Environment Research Council (NERC) and the Food Standards Agency (FSA) under the Environmental Microbiology and Human Health (EMHH) Programme (NE/M010996/1). The authors gratefully acknowledge Dr James Lowther (Centre for Environment, Fisheries and Aquaculture Science; CEFAS) for providing the mengovirus sample. We also thank Gordon Steffen and Dr Nick Barcock (Dŵr Cymru Cyf-Welsh Water Ltd, UK) for facilitating sample collection at the wastewater treatment plants and Dr Julie Webb (Bangor University, UK) for assistance in sampling.

